## Supplementary material for "A rule-based multiscale model of hepatic stellate cell plasticity: critical role of the inactivation loop in fibrosis progression": Sup information

Matthieu Bougueon[1,2], Vincent Legagneux[2], Octave Hazard [4,5], Jeremy Bomo[1,2], Anne Siegel[1], Jérôme Feret[3,4*] Nathalie Théret[1,2*],

**1** Univ Rennes, Inria, CNRS, IRISA, UMR 6074, Rennes, France
**2** Univ Rennes, Inserm, EHESP, Irset, UMR S1085, Rennes, France
**3** Team Antique, Inria, Paris, France
**4** DI-ENS (Inria, ÉNS, CNRS, PSL University), École normale supérieure, Paris, France
**5** École Polytechnique, Palaiseau, France

* *

**Fig S1. Temporal analysis of cells and COL1 in models with or without reversion of iHSC to a quiescent state.**

**Fig S2. Comparison of model predictions and experimental data acquired in models of fibrosis induced by dimethyl_nitrosamine, thioacetamide and a high-fat diet.**

**Table S1. List of the genes used for GSEA analysis.**

| Signature de Rosenthal | | Signature combinée | |
|---|---|---|---|
| Ensembl code | Gene name | Ensembl code | Gene name |
| ENSG00000163739 | CXCL1 | ENSG00000077274 | CAPN6 |
| ENSG00000115380 | EFEMP1 | ENSG00000169245 | CXCL10 |
| ENSG00000148180 | GSN | ENSG00000147257 | GPC3 |
| ENSG00000152583 | SPARCL1 | ENSG00000081041 | CXCL2 |
| ENSG00000112562 | SMOC2 | ENSG00000112799 | LY86 |
| ENSG00000164949 | GEM | ENSG00000163131 | CTSS |
| ENSG00000011465 | DCN | ENSG00000168329 | CX3CR1 |
| ENSG00000177606 | JUN | ENSG00000172602 | RND1 |
| ENSG00000171223 | JUNB | | |
| ENSG00000120738 | EGR1 | | |
| ENSG00000170345 | FOS | | |
| ENSG00000137193 | PIM1 | | |
| ENSG00000142871 | CYR61 | | |
| ENSG00000130522 | JUND | | |
| ENSG00000120708 | TGFBI | | |
| ENSG00000168477 | TNXB | | |
| ENSG00000161544 | CYGB | | |
| ENSG00000272573 | MUSTN1 | | |
| ENSG00000170801 | HTRA3 | | |
| ENSG00000125740 | FOSB | | |
| ENSG00000162545 | CAMK2N1 | | |
| ENSG00000130208 | APOC1 | | |
| ENSG00000171867 | PRNP | | |
| ENSG00000162496 | DHRS3 | | |
| ENSG00000163737 | PF4 | | |
| ENSG00000000971 | CFH | | |
| ENSG00000224389 | C4B | | |
| ENSG00000121207 | LRAT | | |
| ENSG00000144152 | FBLN7 | | |
| ENSG00000143369 | ECM1 | | |
| ENSG00000131620 | ANO1 | | |
| ENSG00000137491 | SLCO2B1 | | |
| ENSG00000146374 | RSPO3 | | |
| ENSG00000165795 | NDRG2 | | |
| ENSG00000147481 | SNTG1 | | |
| ENSG00000133110 | POSTN | | |
| ENSG00000069431 | ABCC9 | | |

### Table S2. Table of parameters.

| Parameter | Symbol/Name | Value | Reference |
|---|---|---|---|
| 1 - Recycling rate | Kr | 4.5 min-1 | [1] |
| 2 - Production rate | Pr | 1.2 min-1 | [1] |
| 3 - Ligand induced degradation rate | Kid | 0.6 min-1 | [1] |
| 4 - Half-life free | TGFB1_half_life | 3 min-1 | Calculated from [2, 3] |
| 5 - Binding time to HSC receptor | TGFB1_binding_HSC | 0.001 min-1 | Calculated from [4] |
| 6 - Binding time to MFB receptor | TGFB1_binding_MFB | 0.004 min-1 | Calculated from [4] |
| 7 - Half-time required for TGF$\beta$1 to induce a signal initiating the activation process | HSC_half_activation | 0.15 hour-1 | Calculated from [5] |
| 8 - Half-time required for TGF$\beta$1 to induce a signal initiating the reactivation process | react_HSC_half_activation | 0.043 hours-1 | [5, 6, 7] |
| 9 - Number of TGFBR by cell | TGFB1_factor | 10 000 | [8, 9, 10] |
| 10 - Saturating quantity of TGF$\beta$1 | TGFB1_input | 60 000/cell | [5] |
| 11 - Number of TGF$\beta$1 input | nb_iteration | 16 | [6] |
| 12 - Time between each TGF$\beta$1 input | interval | 84 hours | [6] |
| 13 - Number of qHSCs at T0 | HSC_cell_density | 10 000 | |
| 14 - Half-life of qHSC | HSC_quiescent_half_life | 75 days-1 | Estimated from biological observations [6] |
| 15 - Half-life of iHSC | iHSC_half_life | 2 years-1 | Estimated from biological observations [6] |
| 16 - Half-life of MFB | MFB_half_life | 3.54 days-1 | Estimated from biological observations [6] |
| 17 - Half-life of react_MFB | react_MFB_half_life | 3.54 days-1 | Estimated from biological observations [6] |
| 18 - Percentage of MFB inactivated | inactivation_percentage | 45% | [6] |
| 19 - Percentage of react_MFB inactivated | inactivation_percentage_react | 5% | Estimated from biological observations [6] |
| 20 - MFB inactivation step half-time | MFB_inact_stage_half_life | 0.75 days-1 | Calculated from [7, 11] |
| 21 - Half life of apop_sene_MFB | apop_sene_MFB_half_life | 3.6 hours-1 | |
| 22 - Proliferating time of aHSC | HSC_doubling_time | 15.5 days-1 | Estimated from biological observations [6] |
| 23 - Proliferating time of react_HSC | react_HSC_doubling_time | 4.43 days-1 | Estimated from biological observations [6, 7, 11] |
| 24 - Proliferating time of MFB | MFB_doubling_time | 31 days-1 | Estimated from biological observations [6, 12, 13] |
| 25 - Proliferating time of react_MFB | react_MFB_doubling_time | 8.86 days-1 | Estimated from biological observations [6, 7, 11] |
| 26 - aHSC activation step half-time | HSC_stage_half_life | 17 hours-1 | Calculated from [14, 13] |
| 27 - MFB differentiation step half-time | MFB_stage_half_life | 17 hours-1 | Calculated from [14, 13] |
| 28 - react_HSC reactivation step half-time | react_HSC_stage_half_life | 4.86 hours-1 | Calculated from [6, 7, 11] |
| 29 - react_MFB differentiation step half-time | react_MFB_stage_half_life | 4.86 hours-1 | Calculated from [6, 7, 11] |
| 30 - % areas in healthy liver | col1_healthy_concentration | 0.5 A.U | [6] |
| 31 - aHSC production of high remodeling COL1 | COL1_synthesis_aHSCs | 0.019 A.U / 12h | Estimated from biological observations [6] |
| 32 - MFB production of high remodeling COL1 | COL1_synthesis_MFB | 0.19 A.U / 12h | Estimated from biological observations [6] |
| 33 - react_HSC production of high remodeling COL1 | COL1_synthesis_react_HSC | 0.0665 A.U / 12h | Estimated from biological observations [6] |
| 34 - react_MFB production of high remodeling COL1 | COL1_synthesis_react_MFB | 0.665 A.U / 12h | Estimated from biological observations [6] |
| 35 - aHSC production of high remodeling COL1 induced by TGF$\beta$1 | COL1_synthesis_aHSC_TGFB1 | 0.19 A.U / 12h | Estimated from biological observations [6] |
| 36 - MFB production of high remodeling COL1 induced by TGF$\beta$1 | COL1_synthesis_MFB_TGFB1 | 0.38 A.U / 12h | Estimated from biological observations [6] |
| 37 - react_HSC production of high remodeling COL1 induced by TGF$\beta$1 | COL1_synthesis_react_HSC_TGFB1 | 0.665 A.U / 12h | Estimated from biological observations [6] |
| 38 - react_MFB production of high remodeling COL1 induced by TGF$\beta$1 | COL1_synthesis_react_MFB_TGFB1 | 1.33 A.U / 12h | Estimated from biological observations [6] |
| 39 - Quantity of degraded high remodeling COL1 and low remodeling COL1 | COL1_deg | 0.0002 A.U | Estimated from biological observations [6] |
| 40 - High remodeling COL1 half life | COL1_remodeling _high_half_life | 17.28 s-1 | Estimated from biological observations [6] |
| 41 - Low remodeling COL1 half life | COL1_remodeling _low_half_life | 15.84 min-1 | Estimated from biological observations [6] |
| 42 - High remodeling COL1 to low remodeling COL1 transition time | COL1_remodeling _high_to_remodeling_low _half_time | 20 days-1 | Estimated from biological observations [6] |
| 43 - Low remodeling COL1 to COL1 High remodeling COL1 transition time | COL1_remodeling _low_to_remodeling_high _half_time | 2.16 min-1 | Estimated from biological observations [6] |

### References


1. Vilar JMG, Jansen R, Sander C. Signal processing in the TGF-$\beta$ superfamily ligand-receptor network. PLoS computational biology. 2006;2(1):e3.

2. Wakefield L, Winokur T, Hollands R, Christopherson K, Levinson A, Sporn M. Recombinant latent TGF-$\beta$1 has a longer plasma half life in rats than active TGF-$\beta$1, and a different tissue distribution. J Clin Invest. 1990;86:1976–1984.

3. Hermonat PL, Li D, Yang B, Mehta JL. Mechanism of action and delivery possibilities for TGF$\beta$1 in the treatment of myocardial ischemia. Cardiovascular research. 2007;74(2):235–243.

4. Friedman SL, Yamasaki G, Wong L. Modulation of transforming growth factor beta receptors of rat lipocytes during the hepatic wound healing response. Enhanced binding and reduced gene expression accompany cellular activation in culture and in vivo. Journal of Biological Chemistry. 1994;269(14):10551–10558.

5. Zi Z, Feng Z, Chapnick DA, Dahl M, Deng D, Klipp E, et al. Quantitative analysis of transient and sustained transforming growth factor-$\beta$ signaling dynamics. Molecular systems biology. 2011;7(1):492.

6. Kisseleva T, Cong M, Paik Y, Scholten D, Jiang C, Benner C, et al. Myofibroblasts revert to an inactive phenotype during regression of liver fibrosis. Proc Natl Acad Sci U S A. 2012;109(24):9448–9453.

7. Massague J, Like B. Cellular receptors for type beta transforming growth factor. Ligand binding and affinity labeling in human and rodent cell lines. Journal of Biological Chemistry. 1985;260(5):2636–2645.

8. Kalter VG, Brody AR. Receptors for Transforming Growth Factor-S (lGF-) on Rat Lung Fibroblasts Have Higher Affinity for IDF-, 81 than for IDF-. Am J Respir Cell Mol BioI. 1991;4:397–407.

9. Lyons RM, Miller DA, Graycar JL, Moses HL, Derynck R. Differential binding of transforming growth factor-$\beta$1,-$\beta$2, and-$\beta$3 by fibroblasts and epithelial cells measured by affinity cross-linking of cell surface receptors. Molecular Endocrinology. 1991;5(12):1887–1896.

10. Troeger JS, Mederacke I, Gwak GY, Dapito DH, Mu X, Hsu CC, et al. Deactivation of hepatic stellate cells during liver fibrosis resolution in mice. Gastroenterology. 2012;143(4):1073–1083.

11. El Taghdouini A, Najimi M, Sancho-Bru P, Sokal E, van Grunsven LA. In vitro reversion of activated primary human hepatic stellate cells. Fibrogenesis & tissue repair. 2015;8(1):1–15.

12. Bachem MG, Meyer D, Melchior R, Sell KM, Gressner AM. Activation of rat liver perisinusoidal lipocytes by transforming growth factors derived from myofibroblastlike cells. A potential mechanism of self perpetuation in liver fibrogenesis. J Clin Invest. 1992;89(1):19–27.

13. Bachem MG, Meyer D, Schäfer W, Riess U, Melchior R, Sell KM, et al. The response of rat liver perisinusoidal lipocytes to polypeptide growth regulator changes with their transdifferentiation into myofibroblast-like cells in culture. Journal of hepatology. 1993;18(1):40–52.

14. Dooley S, Delvoux B, Lahme B, Mangasser-Stephan K, Gressner AM. Modulation of transforming growth factor $\beta$ response and signaling during transdifferentiation of rat hepatic stellate cells to myofibroblasts. Hepatology. 2000;31(5):1094–1106.

15. Sato Y, Yoneda A, Shimizu F, Nishimura M, Shimoyama R, Tashiro Y, et al. Resolution of fibrosis by siRNA HSP47 in vitamin A-coupled liposomes induces regeneration of chronically injured livers. J Gastroenterol Hepatol. 2021;36(12):3418–3428.

16. Abramovitch S, Dahan-Bachar L, Sharvit E, Weisman Y, Ben Tov A, Brazowski E, et al. Vitamin D inhibits proliferation and profibrotic marker expression in hepatic stellate cells and decreases thioacetamide-induced liver fibrosis in rats. Gut. 2011;60(12):1728–1737.

17. Farooq M, Hameed H, Dimanche-Boitrel MT, Piquet-Pellorce C, Samson M, Le Seyec J. Switching to Regular Diet Partially Resolves Liver Fibrosis Induced by High-Fat, High-Cholesterol Diet in Mice. Nutrients. 2022;14(2).

18. Rosenthal SB, Liu X, Ganguly S, Dhar D, Pasillas MP, Ricciardelli E, et al. Heterogeneity of HSCs in a Mouse Model of NASH. Hepatology. 2021;74(2):667–685.
