## Supplementary figures and images for "A rule-based multiscale model of hepatic stellate cell plasticity: critical role of the inactivation loop in fibrosis progression"

### sup FigS1

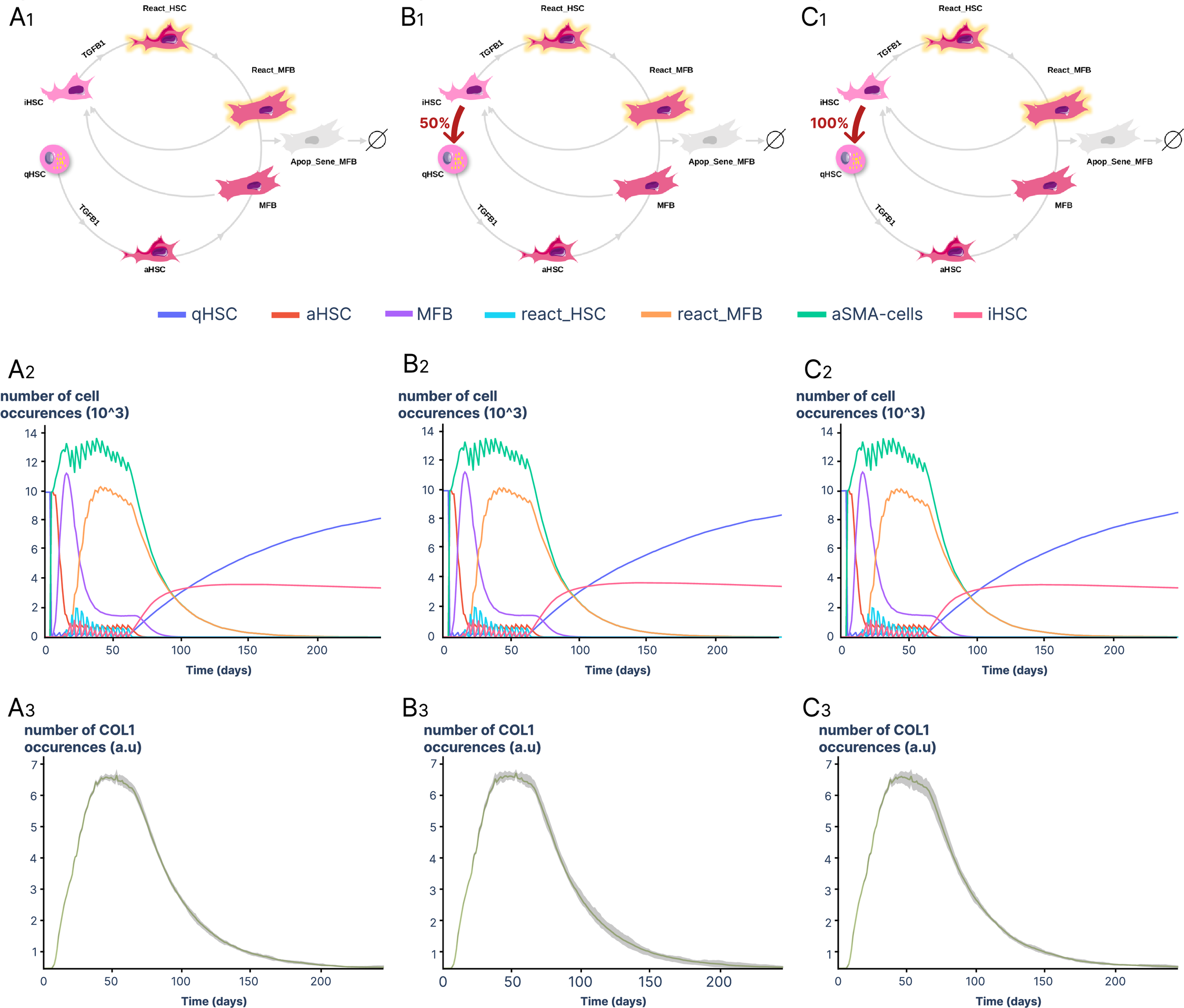

### sup FigS2

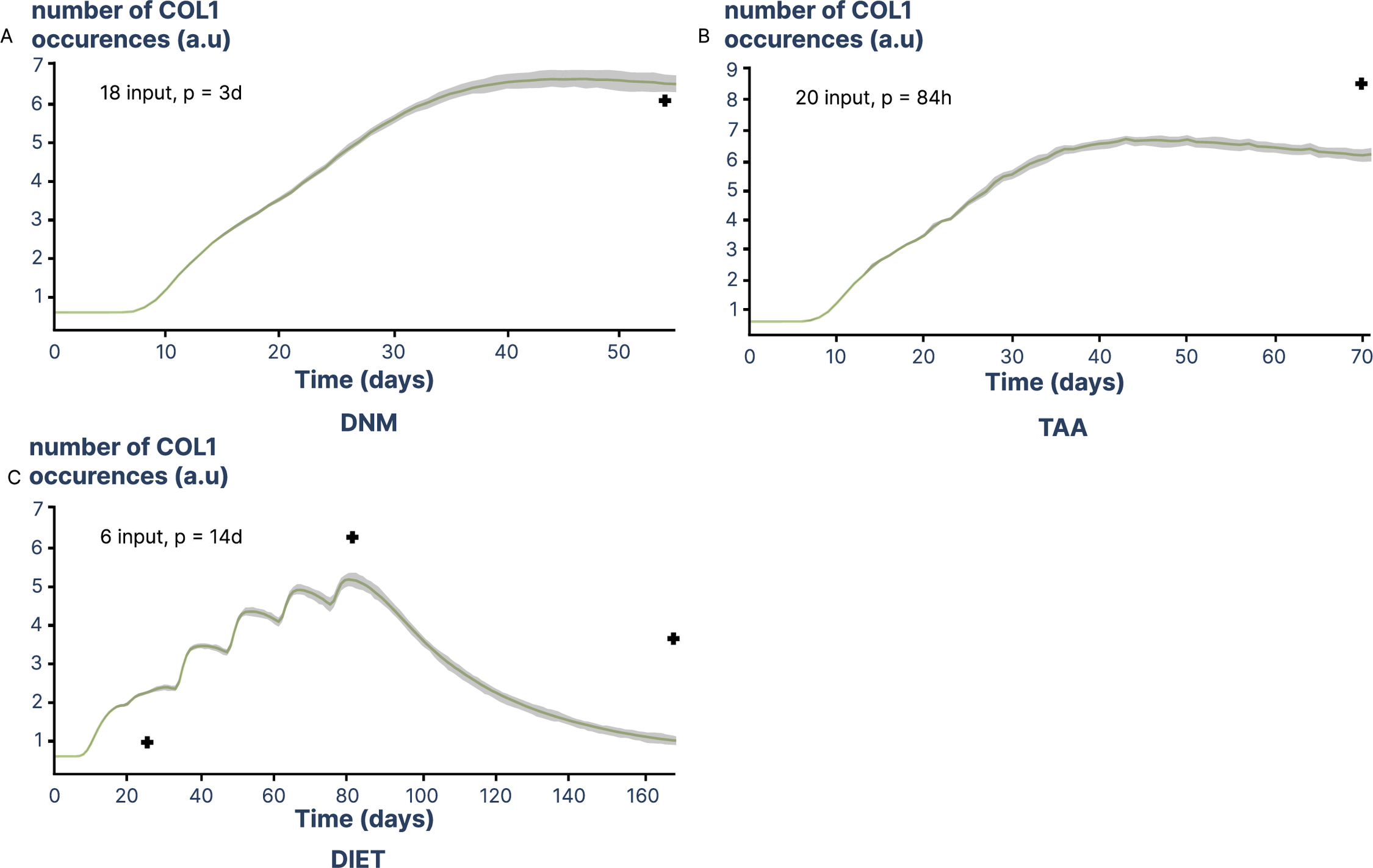
